# Concurrent large-scale brain dynamics during the emotional face matching task and their relation to behavior and mental health

**DOI:** 10.1101/2024.09.21.613739

**Authors:** Cole Korponay, Julia E. Cohen-Gilbert, Poornima Kumar, Nathaniel G. Harnett, Adrian A. Medina, Brent P. Forester, Kerry J. Ressler, Christian F. Beckmann, David G. Harper, Lisa D. Nickerson

**Author notes:** Corresponding author: Lisa D. Nickerson McLean Hospital 115 Mill Street Belmont, MA 02478. These authors contributed equally (Cole Korponay, Julia E. Cohen-Gilbert).

## Abstract

Prior investigations of emotion processing’s neural underpinnings rely on *a priori* models of brain response, obscuring detection of task-relevant neurobiological processes with complex temporal dynamics. To overcome this limitation, we applied unsupervised machine learning to functional magnetic resonance imaging data acquired during the emotional face matching task (EFMT) in healthy young adults from the Human Connectome Project (n=413; n=416 replication). Tensorial independent component analysis showed that the EFMT engages 10 large-scale brain networks – each recruiting visual association cortex in distinct temporal fashions and in tandem with diverse non-visual regions – that collectively recruit 74% of cortex, posterior cerebellum, and amygdala. Despite prominent use of the EFMT to probe negative affect and related psychopathology, EFMT-recruited networks strongly reflected individual differences in cognition but not internalizing/negative affect. Overall, we characterize a richer-than-expected tapestry of concurrent EFMT-recruited brain processes, their diverse activation dynamics, and their relations to task performance and latent mental health phenotypes.

## Introduction

The ability to regulate emotional reactivity amidst the pursuit of task goals is a core pro-social function and is intimately tied to psychiatric well-being and illness^1^. As such, one of the most widely administered and studied tasks using functional magnetic resonance imaging (fMRI) is the emotional face matching task (EFMT)^2,3^. Since its introduction in 2002, the EFMT has been used in over 250 fMRI studies to examine the neurobiological basis of emotion processing, as well as potential links between emotion-related brain activity and neuropsychiatric phenotypes^4^. Early studies using the EFMT primarily focused on group-level, task block-averaged brain activity^2,3^. These studies established robust maps of the brain regions whose activity, for instance, was greater during face-matching trials than shape-matching trials^5^, with the majority of these focused on amygdala activation and amygdala connectivity^6,7^. However, a comprehensive review of such studies shows only sparse associations between these EFMT-based neurobiological phenotypes and psychopathology, despite the examination of multiple disorders and numerous variations of the task^8^.

More recently, two field-wide developments have advanced the focus of inquiry in EFMT studies. The first – motivated by the pursuit of “precision psychiatry” – is a movement away from group-level differences and toward individual-level differences^9^. This has entailed a search for brain-based biomarkers of mental health that are sensitive and specific at the level of individual subjects. However, these efforts have met several challenges^10–12^. One challenge is the larger measurement noise present in individual-level data compared to group-level data for both fMRI and behavioral assays. A second challenge is that, like most fMRI tasks, the EFMT was originally designed to minimize individual variability in brain activation for the sake of maximizing group-level signal^12^. A third challenge is that the space of potential brain-based biomarkers is vast, and it remains unclear which ones (e.g., focal activation magnitudes, connection strengths, etc.) best reflect behavior.

This latter challenge dovetails with the second development in studies of emotion processing and regulation: a shift in focus from focal brain regions to distributed, large-scale brain networks^13–16^. Large-scale brain activation during the EFMT has been analyzed via techniques such as multivoxel pattern analysis^16^, meta-analytic clustering^17^, Bayes factor analysis^14^, and connectome-based predictive modeling^18^, and has expanded the understanding of emotion processing beyond classically implicated structures like the amygdala to brain-wide, interacting and dynamic circuits^8,15,19^. These include networks that sense and integrate emotionally salient environmental signals, such as sensory networks, the ventromedial components of the default mode network (DMN)^18^, and the salience network (SN)^20^, as well as networks implicated in the top-down regulation of attention, cognition and behavior, such as left and right frontoparietal networks (FPN)^21^ and dorsal attention network (DAN)^22^.

However, even these more sophisticated analytical approaches share an underlying similarity with the classic region-of-interest and massively univariate voxel-wise approaches which limits their utility: a reliance on task contrasts. The fundamental input unit to these analyses is a difference score computed between brain activity during task blocks of interest (e.g., emotional faces) and during control blocks (e.g., neutral shapes). This task contrast method has proven powerful in identifying brain regions and networks whose activity significantly changes during blocks of interest relative to baseline. However, this top-down approach artificially constrains the identification of potentially relevant brain activation patterns to those in the experimenters’ predefined model of how the measured fMRI signal is modulated by the task. Moreover, the reliance of these approaches on block-averaged brain activity obscures fine-grained temporal dynamics that might occur within and between blocks.

An alternative approach would involve a bottom-up, data-driven search for brain activity whose time course aligns with one or more aspects of the task stimulation time course, and which retains fine-grained temporal information. A well-validated method that can extract this kind of information from task fMRI data is tensor independent component analysis (tICA^23^), an unsupervised learning technique that can detect and trace the moment-by-moment activity of large-scale, distributed brain networks during task performance. However, previous applications of tICA to task fMRI have focused primarily on assessing subject differences in^24^ or methodological optimizations yielded by^25^ tICA-identified networks, rather than on deeply characterizing the neurobiological features and functions of brain networks during task performance. Given that the EFMT has been included in multiple large-scale neuroimaging datasets, including the Human Connectome Project (HCP), UK Biobank, and in several of the Connectomes Related to Human Disease as a probe of the NIMH Research Domain Criteria (RDoC)-based Negative Valence System^26^, and is used frequently to assess individual differences in negative affect, applying tICA to the EFMT stands to significantly deepen the understanding of brain network dynamics and their relation to cognitive and psychological factors during this important task.

In the current study, we applied tICA to EFMT fMRI data from the HCP Young Adult sample to detect and trace the activity of large-scale brain activation patterns during different temporal sub-domains of the task (e.g., during the onset of emotional face blocks, throughout emotional face blocks, during the offset of blocks, in between emotional face blocks and neutral shape blocks, etc.). Thus, we used tICA to identify – and characterize at high spatiotemporal resolution – EFMT-recruited neuronal processes that may be missed or obscured by the constraints of contrast-focused analyses^5^. We also used tICA to generate individual subject loadings for each identified brain network, to assess how individual differences in network activity relate to individual differences in task performance, as well as cognitive/behavioral and mental health factors.

## Results

### The EFMT Reproducibly Recruits 10 Large-Scale Networks

The HCP sample was split into two groups, Group 1 (n=413) for network identification and a replication sample (Group 2; n=416), with all members of a family included in the same group to prevent family structure-related data leakage across the two groups. Two runs from each participant (with left-right and right-left phase encoding, hereafter referred to as LR/RL) were analyzed separately. tICA identified 10 spatiotemporally distinct large-scale brain networks recruited by the EFMT in Group 1’s first run (i.e., LR run) (**Fig. 1**), which were robustly replicated in Group 2’s LR EFMT run (**Table 1**). Two of the ten networks did not replicate in the second EFMT runs for each group (i.e., the RL runs), suggesting that the engagement of some networks may be sensitive to task practice, habituation or fatigue effects within the scanning session. Given differences between the first (LR) and second (RL) runs, we focused subsequent analyses on the first LR run from Group 1, with replications in the first LR run from Group 2.

**Figure 1.**
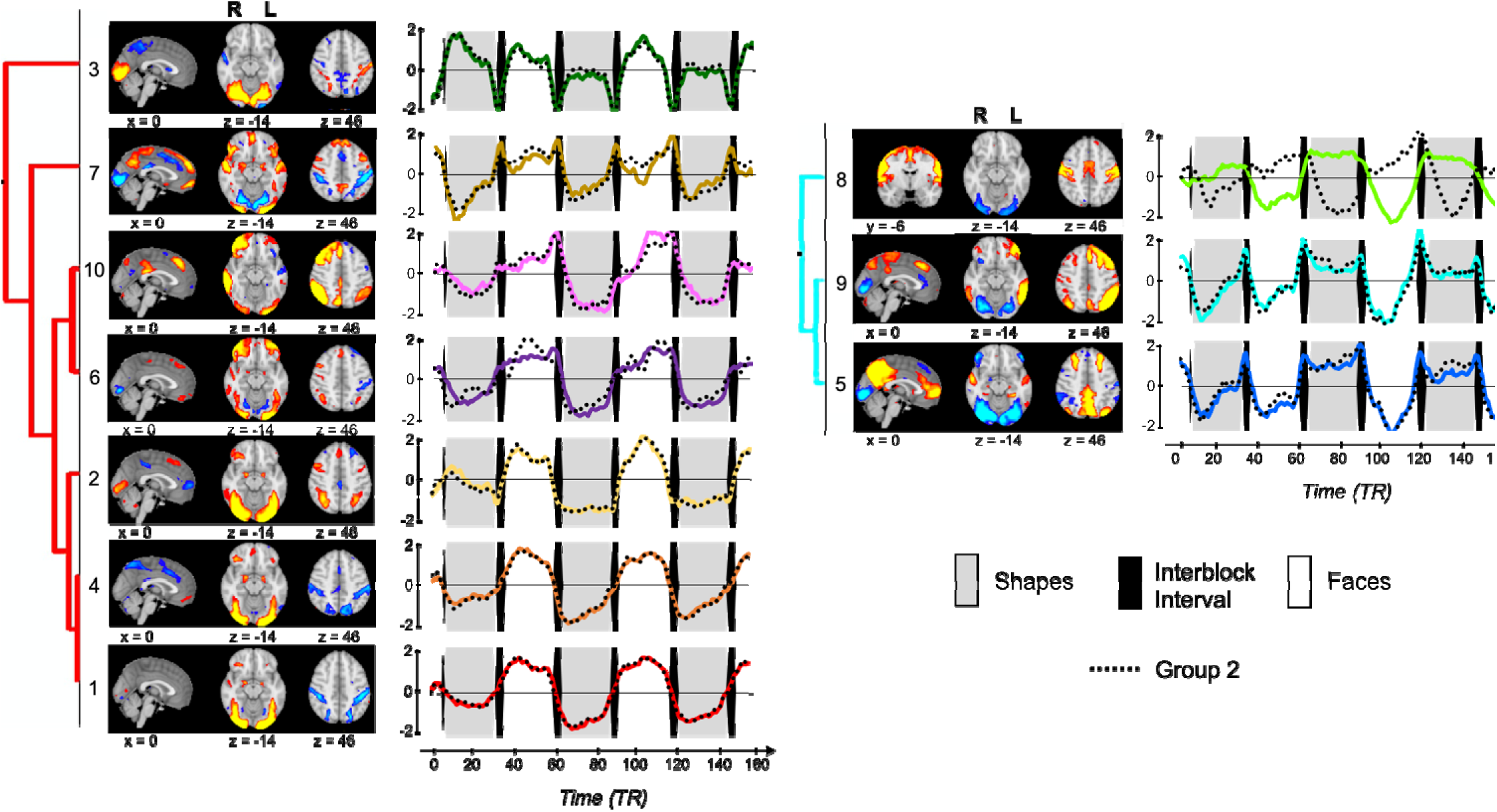
Ten large-scale brain networks engaged during the EFMT. The spatial maps and temporal dynamics of EFMT-recruited networks identified by tICA. Hierarchical clustering distinguished networks preferentially activated for faces > shapes (left panel) and for shapes > faces (right panel) in analyses conducted post-hoc on the subjects’ loadings identified by tICA. Colored lines show the temporal courses for networks identified in Group 1 LR, and dotted lines show the temporal courses of the corresponding networks identified in Group 2 LR.

**Table 1.** Reproducibility of EFMT-recruited networks across groups and runs. Spatial map correlations and temporal course correlations between the Group 1 LR reference networks and the closest corresponding networks in Group 2 LR, Group 1 RL, and Group 2 RL. Networks without a corresponding match (*r*<0.50 for both spatial and temporal correlation) are left blank. *The EFMT Recruits Most of Cortex and Circumscribed Non-Cortical Areas*

| Group 1 LR Network | | Spatial $r$ | Temporal $r$ | | Spatial $r$ | Temporal $r$ | | Spatial $r$ | Temporal $r$ |
| --- | --- | --- | --- | --- | --- | --- | --- | --- | --- |
| 1 | Group 2 LR Network Match: | 0.96 | 0.99 | Group 1 RL Network Match: | 0.94 | 0.98 | Group 2 RL Network Match: | 0.92 | 0.97 |
| 2 |  | 0.93 | 0.99 |  | - | - |  | 0.77 | 0.97 |
| 3 |  | 0.95 | 0.98 |  | 0.88 | 0.95 |  | 0.85 | 0.95 |
| 4 |  | 0.96 | 0.99 |  | 0.84 | 0.93 |  | 0.56 | 0.85 |
| 5 |  | 0.77 | 0.95 |  | 0.65 | 0.73 |  | 0.68 | 0.82 |
| 6 |  | 0.48 | 0.90 |  | 0.71 | 0.95 |  | 0.66 | 0.95 |
| 7 |  | 0.78 | 0.92 |  | 0.69 | 0.75 |  | 0.83 | 0.90 |
| 8 |  | 0.76 | -0.53 |  | 0.71 | 0.80 |  | - | - |
| 9 |  | 0.91 | 0.97 |  | - | - |  | - | - |
| 10 |  | 0.88 | 0.95 |  | 0.93 | 0.86 |  | 0.94 | -0.29 |

Spatially, all ten networks included the fusiform gyrus and higher-level visual cortex as nodes (**Fig. 2**). Nine of the ten included the right pars triangularis of the inferior frontal gyrus (BA45), and seven of the ten included primary somatosensory cortex (BA1). Collectively, 74% of cortex was recruited as a node in one or more of the large-scale networks during the EFMT. Notably absent as a node in any of the ten networks was the subgenual cingulate cortex as defined by Mayberg and colleagues^27^, a brain region strongly implicated in negative affect regulation and depression^28,29^. Outside of cortex, seven of the ten networks included the posterior cerebellum as a node, and six of ten included the amygdala. Other than the amygdala, subcortical recruitment was sparse, with parts of striatum, thalamus and hippocampus involved in only one network each.

**Figure 2.**
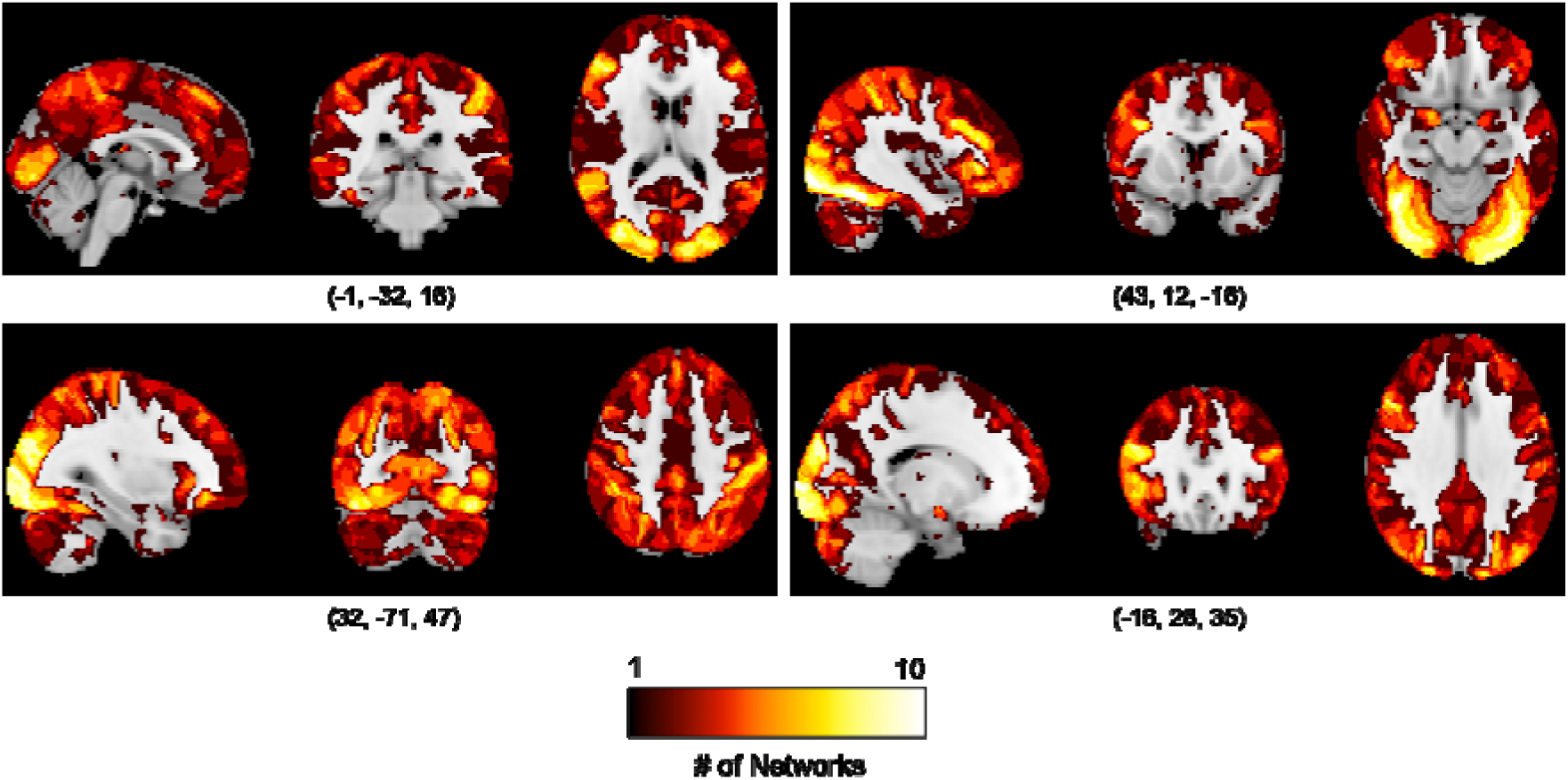
Spatial overlap of brain networks recruited during EFMT. A global view of brain involvement in the EFMT. The positive and negative components of each network from Group 1 LR were thresholded at p=0.5 (mixture-modeling), binarized, combined, and summed. Unshaded (i.e., greyscale) regions did not appear as nodes in any of the EFMT-recruited networks. This network overlap map strongly replicated in Group 2 LR (*r*=.91, *p*<.001).

To inform the potential role of each task network in task performance, spatial overlap between the EFMT networks and canonical brain networks that have been previously reported was also assessed. Overlap between the positive and negative components of each network and each of 17 canonical large-scale networks defined in the Gordon et al. parcellation^30^ was computed using the network correspondence toolbox (NCT)^31^. Results of this analysis are presented in **Fig. 3** and described further in the in-depth network profiling below.

**Figure 3.**
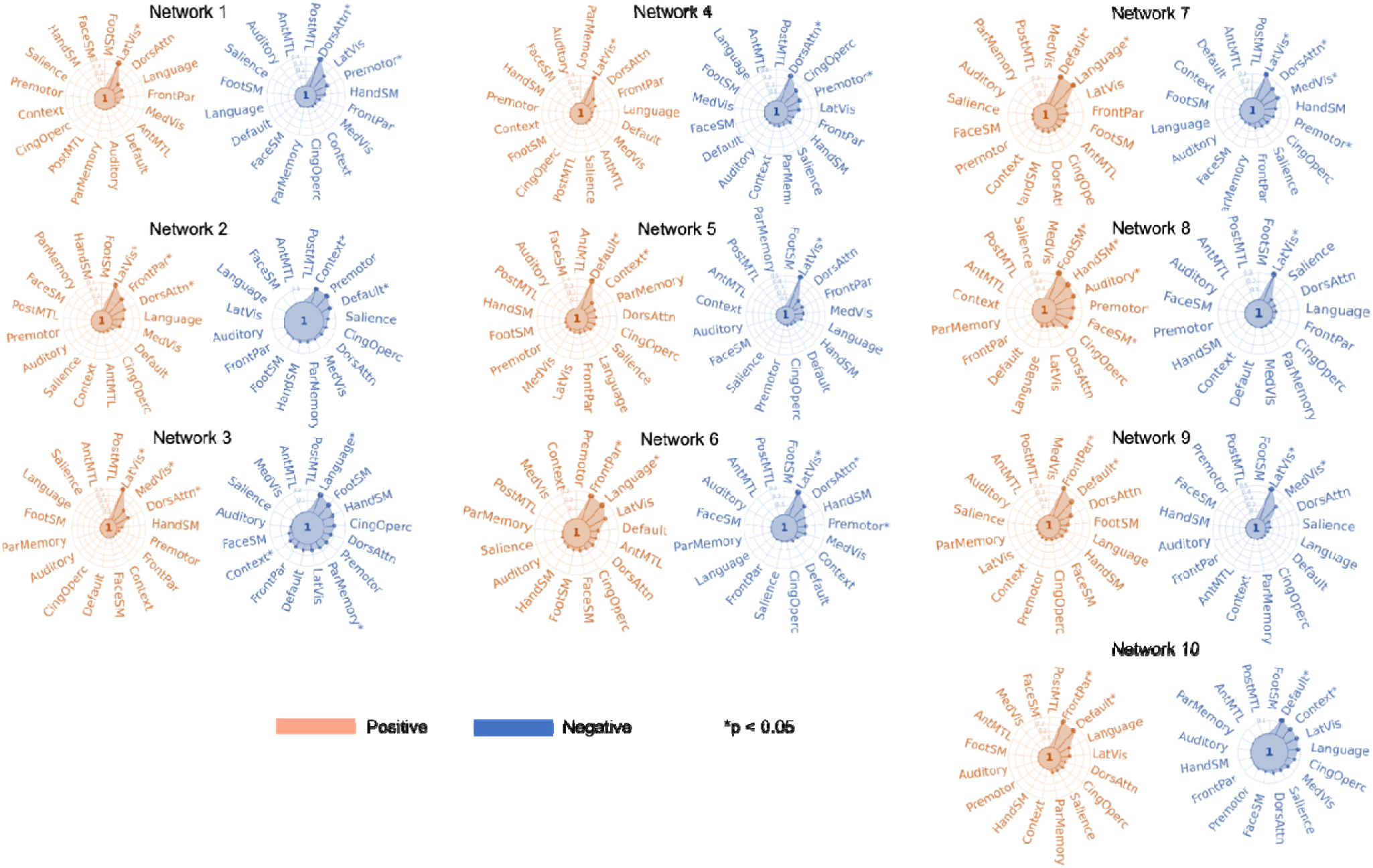
Overlap of EFMT-Networks with Canonical Networks. Plots display Dice coefficients generated by the NCT^31^ quantifying the extent of spatial overlap between the positive (red) and negative (blue) components of each EFMT-recruited network (Group 1 LR) and each of 17 canonical large-scale networks defined in the Gordon et al. parcellation^30^. Radar charts show the overlapping atlas networks with greatest to lowest Dice coefficients clockwise around the graph. Findings were consistent in Group 2 LR. Networks include: default mode (Default); medial and lateral visual (MedVis, LatVis); cingulo-opercular (CingOperc); salience; frontoparietal (FrontPar); dorsal attention (DorsAttn); hand, face, and leg somatomotor (HandSM, FaceSM, FootSM); auditory; premotor; parietal memory (ParMemory); contextual association (Context); and anterior and posterior medial temporal networks (AntMTL, PostMTL).

### EFMT-Recruited Networks Have Diverse Temporal Dynamics

Hierarchical clustering of the network time course correlation matrix (**Fig. S1**) revealed two broad types of networks: those preferentially activated during emotional faces trial blocks (**Fig. 1**, left panel) and those preferentially activated during neutral shapes trial blocks (**Fig. 1**, right panel). This clustering is further reflected in the correspondence between the network time courses and EFMT stimulation time courses, identified by the *post hoc* fitting of network timeseries data with a standard block design general linear model (GLM; **Table 2**). At deeper levels of the clustering hierarchy, fine-grained temporal dynamics revealed several further distinctions. For instance, while the engagement of networks 1, 2 and 4 peaked in the middle of emotional face blocks and began tapering off at the end of these blocks, the engagement of networks 6 and 10 ramped up across emotional faces trials and peaked at the end. Meanwhile, network 7 engagement peaked at the onset and offset of emotional faces trials. Dashed lines in the time courses of the networks showing the time courses in the replication sample show strong replication, although it is noted that the time course of network 8 in the replication sample has a negative correlation with the time course from Group 1, likely a result from arbitrary sign flips between modes in the tensor ICA (e.g., spatial maps and time courses can both be multiplied by -1, time courses and subject loadings by -1, or spatial maps and subject loadings by -1 as equivalent solutions to the tensor decomposition).

**Table 2.** Correspondence of brain network time courses with EFMT stimulation time courses. Network time courses identified by tICA were fit *post hoc* with a standard block design general linear model, showing that our data-driven approach identifies networks modulated by the task. Refer to **Fig. S4** for visualization of the stimulation time courses.

|  | 1 | 4 | 2 | 6 | 10 | 7 | 3 | 5 | 9 | 8 |
| --- | --- | --- | --- | --- | --- | --- | --- | --- | --- | --- |
| Faces > Shapes | 20.85 | 20.6 | 15.97 | 20.38 | 13.81 | 10.86 | 8.19 |  |  |  |
| Shapes > Faces |  |  |  |  |  |  |  | 12.49 | 8.6 | 17.15 |
| Faces, Positive | 10*** | 12.66*** | 12.12*** |  |  |  | 13.18*** |  |  |  |
| Faces, Negative |  |  |  |  |  | 8.05*** |  | 12.79*** | 10.78*** | 11.4*** |
| Shapes, Positive |  |  |  |  |  |  | 11.08*** |  |  |  |
| Shapes, Negative | 12.25*** | 8.59*** |  | 16.42*** | 10.25*** | 12.96*** |  | 7.72*** | 7.54*** |  |
| Faces Derivative, Positive |  |  |  |  |  |  | 4.03* |  |  |  |
| Faces Derivative, Negative |  | 5.14*** |  | 9.15*** | 6.81*** | 9.48*** |  |  | 6.93*** |  |
| Shapes Derivative, Positive |  |  | 3.87* |  |  |  | 7.94*** |  |  |  |
| Shapes Derivative, Negative | 4.40** | 7.12*** |  | 9.27*** |  | 12.27*** |  | 6.65*** | 6.12*** |  |

\*p < .0001, \*\*p < .00001, \*\*\*p < .00000

### Individual Differences in EFMT-Recruited Networks and Task Performance

All individual differences analyses controlled for age, sex and education and corrected for covariance in family structure (i.e., family members with non-independent data) using permutation analysis of linear models (PALM)^32^ with multi-level exchangeability blocks^33^. Three participants (two from Group 1 and one from Group 2) were removed from analyses examining brain-behavior associations due to low overall EFMT accuracy (<70%).

Given very high levels of accuracy on both faces and shapes conditions (**Fig. S2**), leading to potential ceiling effects for these metrics, we operationalized EFMT performance by reaction time slowing on emotional faces trials relative to shapes trials, i.e., Rt_faces_ – Rt_shapes_, indexing “emotion interference” (**Fig. S3**). The emotional face condition elicits an emotional reaction from participants that requires early attentional screening and downregulation of salient yet extraneous emotional information, to focus on the task-relevant aspects of the stimulus (i.e., matching the face identity). The difference in reaction time on emotional face trials versus neutral shapes trials thus gauges the extent of this “emotion interference” process. Individual differences in this measure of emotion interference were significantly associated with individual differences in the engagement of four networks at p<.005 (Bonferroni-corrected for multiple comparisons), and of seven networks at the uncorrected level (p<0.05) in Group 1. These significant associations were all in the positive direction, with the exception of Network 7. Relationships for Networks 2 and 10 were the most robust, replicating at the Bonferroni-corrected level in Group 2 (**Fig. 4**). The negative association observed for Network 7 did not replicate in Group 2.

**Figure 4.**
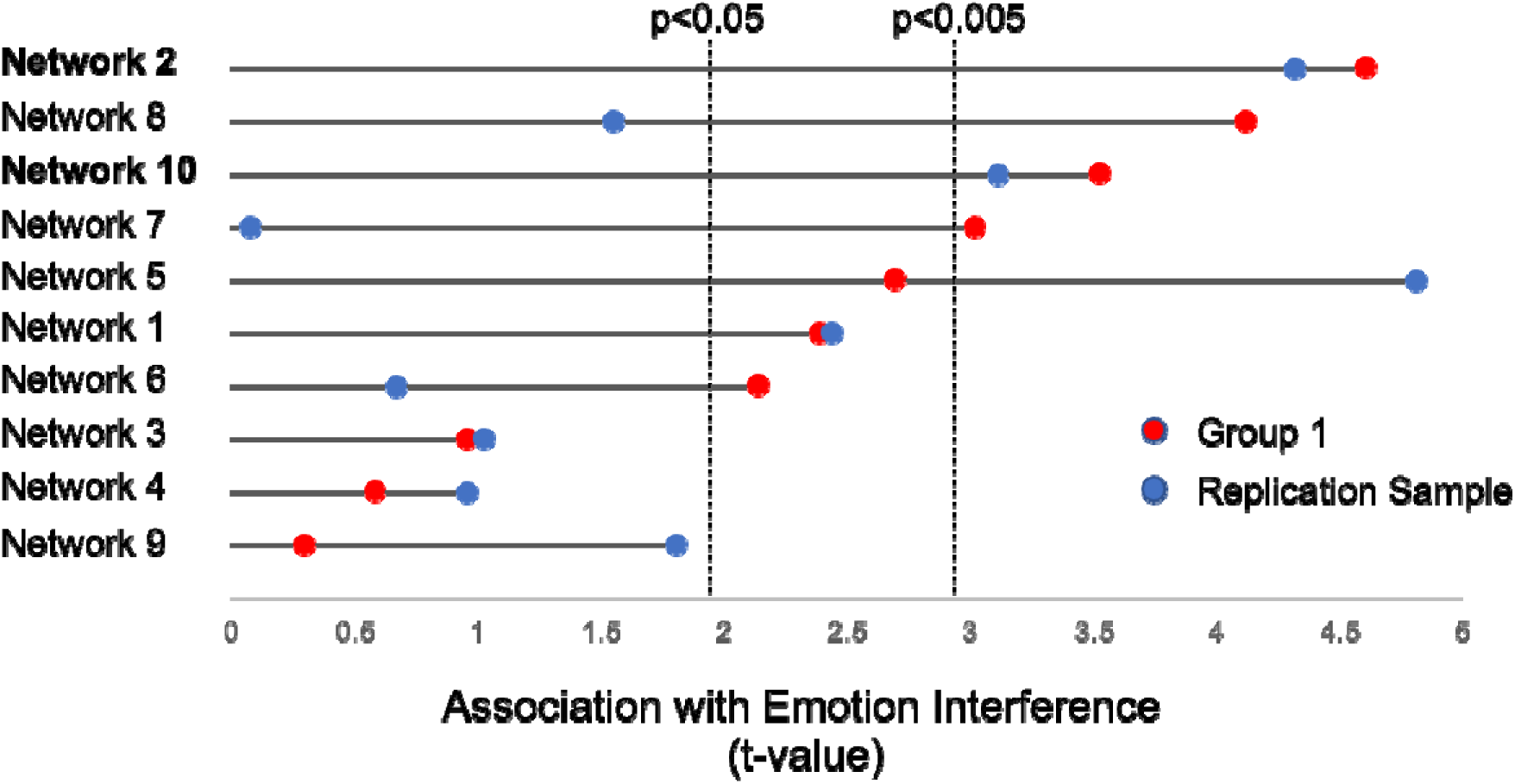
Individual differences in brain network engagement and EFMT task performance. For Group 1 and for the replication sample (Group 2), the relationship between subject loadings on each large-scale network and reaction time slowing on emotional face trials (i.e., emotion interference), controlling for age, sex and education, and correcting for covariance in family structure using permutation analysis of linear models (PALM). Networks with significant relationships in both groups that are below the Bonferroni threshold for multiple comparisons correction (p<0.005) are bolded.

### Individual Differences in EFMT-Recruited Networks and Cognitive/Mental Health Factors

To examine relationships between EFMT networks and latent psychobehavioral phenotypes, we first implemented hierarchical factor analysis on 87 variables from across the HCP’s assessments of alertness, cognition, emotion, motor function, personality, sensory function, psychiatric and life function, substance use and in-scanner tasks^34^. We used the five-factor solution characterized by Schottner and colleagues^34^, comprised of well-being/positive affect, internalizing/negative affect, processing speed, cognition, and substance use. Individual differences in the engagement of five different networks were reproducibly associated with individual differences in cognition across both groups (**Fig. 5**). However, none of the networks were associated with individual differences in positive affect/well-being, negative affect/internalizing, processing speed, or substance use. Networks 4 and 9 were not associated with individual differences in task performance or overall cognition in either group, despite these networks’ strong alignment with multiple subdomains of the task (**Fig. 1**; **Table 2**).

**Figure 5.**
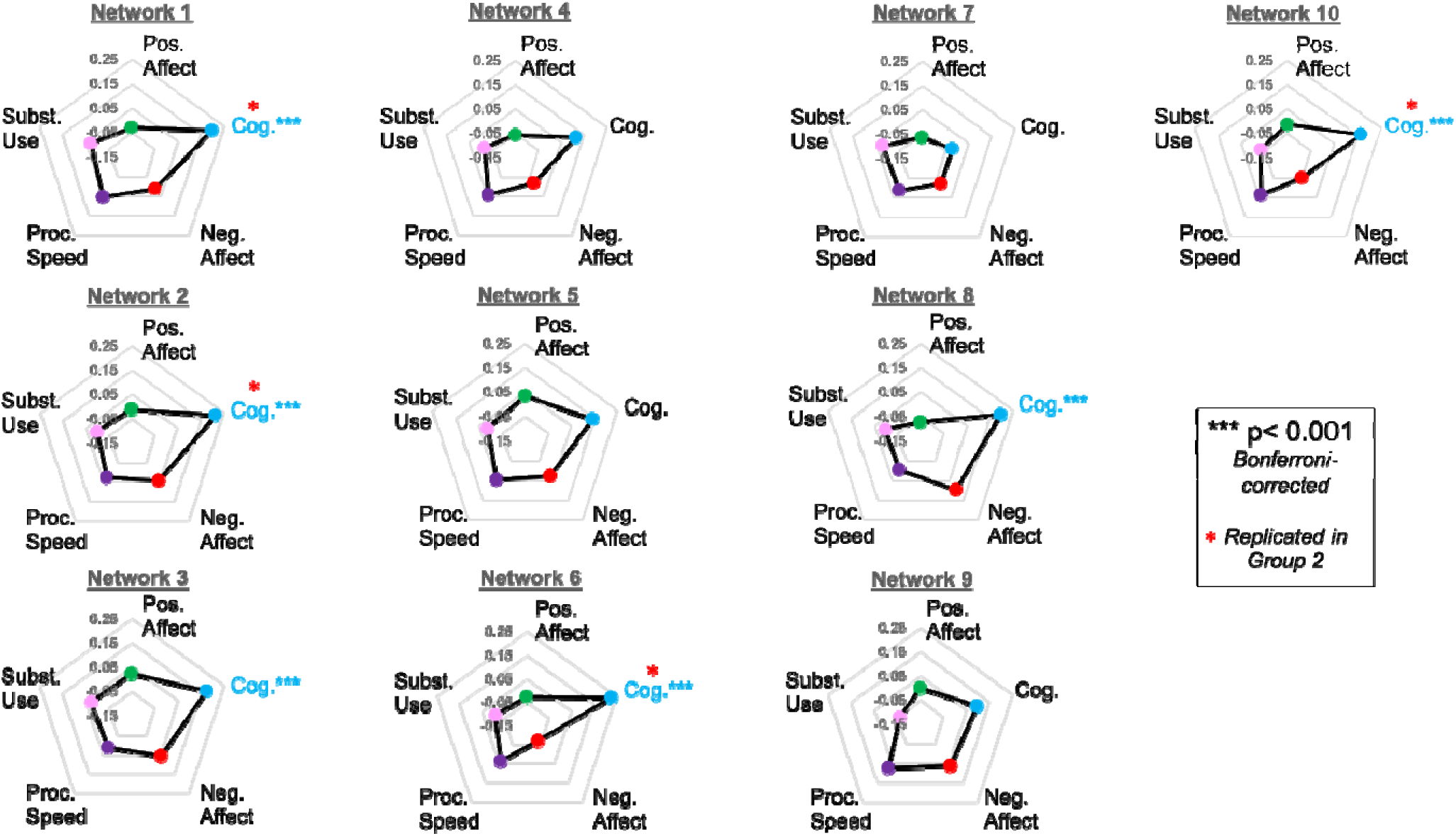
Individual differences in brain network engagement and cognitive/mental health factors. Group 1 relationships between subject loadings on each large-scale network and five facets of behavior derived by factor analysis from 87 HCP-YA behavioral and neuropsychiatric assays: positive affect/well-being, cognition, negative affect/internalizing, processing speed, and substance use. Analyses control for age, sex and education, and correct for covariance in family structure using permutation analysis of linear models (PALM). Significant relationships below the Bonferroni threshold for multiple comparisons correction (p<0.001) are denoted.

### Network Profiling in Depth

#### Networks 1, 2 and 4

Networks 1, 2 and 4 showed peak activation in the middle of emotional faces blocks (**Fig. 1**). The positive spatial components of all three networks (i.e., those activated during face blocks) involved the lateral visual network (**Fig. 3**), including visual association areas such as bilateral inferior and middle occipital cortex and bilateral lingual and fusiform gyri. The amygdala and ventrolateral prefrontal cortex (VLPFC) were also activated during face blocks in all three networks.

The negative spatial components of Networks 1 and 4 (i.e., those suppressed during face blocks) involved the dorsal attention network (DAN) and premotor network. In contrast, the DAN was involved in the positive component of Network 2 (**Fig. 3**), and the activated DAN regions were interdigitated^35^ between the DAN areas suppressed in Networks 1 and 4. Network 2 also differed spatially from Networks 1 and 4 via activation of dorsal brain regions implicated in visual processing and attention (i.e., superior parietal, dorsolateral and dorsomedial prefrontal cortices), more extensive activation of VLPFC and VMPFC, and suppression of the DMN and contextual association network^36^. The contextual association network from the Gordon et al. parcellation^30^ corresponds most closely with Default Network C from the Yeo et al. 17-network parcellation^31,35^ and the medial temporal DMN subsystem defined by Andrews-Hanna and collegues^37^.

Temporally, Network 2 activation occurred slightly after Networks 1 and 4 activation during the onset of face blocks (**Fig. S5**). Furthermore, while Networks 1, 2 and 4 were all positively associated with the faces blocks, only Networks 1 and 4 were significantly negatively associated with shapes blocks (Table 2). Inspection of the time courses indicates this may reflect a lack of suppression of Network 2 during the first shapes block (**Fig. 1**).

Individual differences in the activation of Networks 1 and 2 were significantly associated (p<.001) with individual differences in cognition, with both findings replicating in Group 2; Network 4 was not reproducibly associated with any of the psychobehavioral factors (**Fig. 5**). Amongst these three networks, only individual differences in the activation of Network 2 were significantly positively associated (p<.005) with individual differences in emotion interference (RT_faces_ – RT_shapes_), which was replicated in Group 2 (**Fig. 4**).

#### Networks 6 and 10

Networks 6 and 10 showed peak activation at the end of emotional faces blocks (**Fig. 1**), wherein activation gradually increased across the block. As such, the significant faces>shapes contrast for these networks was driven by significant suppression during shapes blocks, rather than by significant activation during face blocks.

Despite their similar time courses (**Fig. S6**), Networks 6 and 10 were mostly spatially distinct. The positive component of both networks included the frontoparietal network (FPN) (**Fig. 3**) and parts of visual association cortex and right VMPFC. However, for Network 6, the positive component also included the language network and the negative component included the DAN, lateral visual and premotor networks. Meanwhile, both the positive and negative components of Network 10 showed significant overlap with the DMN, and the negative component also included the contextual association network.

Individual differences in the activation of both Networks 6 and 10 were significantly associated (*p*<.001) with individual differences in cognition, with both findings replicating in Group 2 (**Fig. 5**). However, only individual differences in the activation of Network 10 were significantly positively associated (*p*<.005) with individual differences in emotion interference (RTfaces – RTshapes); this was replicated in Group 2 (**Fig. 4**).

#### Networks 3 and 7

Network 7 showed peak activation during interblock intervals, while Network 3 showed peak deactivation during interblock intervals. Both Networks 3 and 7 were positively associated with the Faces > Shapes contrast. However, when each block condition was analyzed separately, Network 3 was positively associated with both faces and shapes conditions, while Network 7 was associated negatively with both conditions (**Table 2**). Closer examination of these networks’ time courses (**Fig. S7**) revealed that these relationships change over the course of the run: Network 3 is activated during the first shapes block but not during subsequent shapes blocks, while the suppression of Network 7 decreases with each subsequent shapes block (**Fig. 1**).

The positive and negative components of these networks’ spatial maps also reflect this spatially overlapping but oppositional theme. The NCT analysis showed significant overlap between both lateral and medial visual networks and the positive component of Network 3, but the negative component of Network 7. Likewise, the NCT analysis showed significant overlap between both language network and DAN and Networks 3 and 7 but opposite, e.g., negative and positive components, respectively. However, the negative component Network 3 also overlapped significantly with the parietal memory network while the positive component of Network 7 comprised parts of DMN and the negative component of Network 7 included premotor areas.

Individual differences in the activation of Network 3, but not Network 7, were significantly associated (*p*<.001) with individual differences in cognition, with a trending relationship (*p*=.006) in Group 2 (**Fig. 5**). Conversely, individual differences in the activation of Network 7, but not Network 3, were significantly associated with individual differences in emotion interference (RT_faces_ – RT_shapes_), in Group 1; however, this relationship did not replicate in Group 2 (**Fig. 4**).

#### Networks 5 and 9

Networks 5 and 9 showed peak activation during the interblock intervals (**Fig. 1**) and both were suppressed during face blocks. While both were also significantly negatively associated with the shapes contrast, detailed examination of the network time courses (**Fig. S8**) revealed that suppression only occurred during the first shapes block, and that the suppression decreased over the course of this block. For Network 9, this pattern of tapering suppression is also observed for faces blocks.

The positive components of both networks included the DMN, and the negative components of both networks included the lateral visual network. The positive component of Network 5 also included the contextual association network, while the positive component of Network 9 included the left frontoparietal network.

Individual differences in the activation of these networks were not significantly related to individual differences in any of the five psychobehavioral factors (**Fig. 5**). However, individual differences in the activation of Network 5 were significantly positively associated (p<.005) with individual differences in emotion interference in the replication sample with the relationship approaching significance in Group 1 (**Fig. 4**).

#### Network 8

Network 8 activation peaked during shapes blocks; its only significant contrast association was a negative relationship with face blocks. The positive component of Network 8 comprised mainly sensorimotor regions while the negative component comprised lateral visual cortex. Replication of this network’s time course was not as strong as for other network time courses (**Table 1**). Individual differences in the activation of Network 8 were significantly associated (*p*<.001) with individual differences in cognition (**Fig. 5**) and with individual differences in task performance (*p*<.005) (**Fig. 4**), although neither was replicated.

### Comparisons to Amygdala Connectivity and Activation during the EFMT

A central focus of prior research on the EFMT’s clinical utility has been on amygdala activation and connectivity^8^. To compare the task performance and psychobehavioral information reflected in amygdala connectivity and activation to that in the presently identified large-scale networks, we used dual regression^38^ to compute functional connectivity between the left and right centromedial amygdala, basolateral amygdala, and superficial amygdala and each of the 10 large-scale networks. We also extracted the task activation spatial maps of regression-weights^5^ from the faces > shapes contrast computed using a standard voxel-wise general linear model that were released by the HCP. From these maps, we calculated the average regression weight in each of the six amygdala regions of interest for each participant. Despite some relationships between amygdala connectivity and activation and task performance and cognitive/mental health domains, none survived Bonferroni correction for multiple comparisons, and effect sizes were considerably smaller compared to the brain-behavior findings using the large-scale networks. As such, tICA-derived networks appear to be a more robust neural correlate of EFMT task performance and general cognition than amygdala connectivity or activation.

## Discussion

The present study reveals the utility of going beyond model-driven -GLM or multiple regression - contrast-based analyses of task fMRI data. Voxel-wise GLM analyses applies the same *a priori* specified model of the brain’s temporal response at every voxel, with resulting contrast maps (e.g., faces > shapes) reflecting an aggregate map comprised of any voxel significantly activated by the task, according to the *a priori* model. This map may be a mix of multiple networks. In addition, brain regions and networks with temporal responses to the task that differ from the *a priori* time courses in the model will not be identified. Here, using tICA, we demonstrate that EFMT-recruited brain networks with greater activity during face matching relative to shapes (i.e., Networks 1, 2, 3, 4, 6, 7, and 10) have diverse temporal activation patterns that may reflect distinct aspects of task performance. For instance, interindividual variance in activation of Network 10, which ramped up during face blocks, was significantly associated with emotion interference, whereas interindividual variance in Network 1 activation, which remained steadily high during face blocks, was not. Several networks also displayed changes in their activation patterns across the course of the task run. For instance, Network 3 displayed strong activation during the first shape block, but little to no activation during subsequent shape blocks. These within-block and within-run network activation changes (**Fig. 1**) identified using the tICA data-driven approach are not captured by standard voxel-wise GLM analyses, that model each task condition as a block convolved with a hemodynamic response function, and may reflect overlooked (or difficult to model) effects of practice, habituation, or fatigue. Overall, tICA’s use of fine-grained temporal information identified a richer-than-expected tapestry of concurrent neurobiological processes recruited by the EFMT, distinguishing brain networks with diverse activation patterns and with distinct relations to task performance and general cognition.

The ten EFMT-recruited networks identified by tICA involve the interaction of visual association cortex with different sets of non-visual brain regions in distinct temporal fashions. These diverse interactions are supported by the multifaceted anatomical connections between visual cortex and the rest of the brain. Kravitz et al., 2013^39^ detail the ventral visual network’s projections to at least six distinct areas that support different functions, including an occipitotemporal-amygdala pathway involved in detecting and processing emotionally salient stimuli and an occipitotemporal-VLPFC pathway supporting object working memory. Networks 2 and 6 show extensive recruitment of both the occipitotemporal-VLPFC pathway and the occipitotemporal-amygdala pathway. Network 2 remains active throughout face blocks, suggesting its engagement of working memory processes that support maintenance of the task rule set despite the presence of salient emotional stimuli, supported by DAN-mediated spatial executive attention. On the other hand, Network 6 activity increases over the course of each face block, and as such may support the focus of visual attention as the interference load created by the emotional faces accrues across the block. Network 10 also increases in activation over the course of the face blocks, but this network’s engagement of the right-lateralized FPN suggests a role in top-down executive control over emotional reactivity.

Notably, Network 2 (visual association + DAN + VLPFC) along with Network 10 (visual association + left FPN) were both significantly and reproducibly associated with individual differences in emotion interference during the EFMT as well as with general cognition. Both networks were preferentially activated during face blocks but had distinct time courses, with Network 2 activity peaking in the middle of face blocks and Network 10 activity peaking at the end of blocks. This points to the two networks’ distinct roles and timing in the emotion regulation process. Network 5 (DMN + contextual association network) was also significantly associated with emotion interference, but in contrast to Networks 2 and 10, it was preferentially activated during interblock intervals and suppressed during face blocks, which aligns with the DMN’s role as a task-negative network. This suggests that the capacity to suppress internally-focused thoughts during active periods of the task, and/or the ability to prepare for upcoming blocks during task rest periods, can also impact task performance.

Even though individual differences in the recruitment of the remaining seven networks were not robustly associated with variability in emotion interference, they almost certainly play roles in supporting other cognitive aspects of task engagement and completion, as supported by the significant relations between several of these networks and general cognition (**Fig. 5**). The emotional face matching condition and shape matching (control) condition differ in several aspects besides emotion content that require engagement of multiple processes. First, the face stimuli are considerably more complex than the ovals used in the shape-matching condition, requiring modulation of focus and cognitive effort. Further, visual processing of human faces by other humans employs specialized brain mechanisms that enable rapid holistic assessment of facial identity and emotion^40^, requiring recruitment of distinct visual processing streams by the two conditions. Finally, the emotional face condition requires early attentional screening and later downregulation of salient yet extraneous emotional information to focus on the task-relevant aspects of the stimulus (i.e., matching the face identity). Each of these cognitive aspects of the task likely follow distinct time courses and are differentially impacted by practice, fatigue, repetition priming, and habituation.

Networks 1, 2 and 4 appear to be engaged in one or more of these distinct cognitive components of the task. Networks 1 and 4 show increased engagement of the ventral visual stream, amygdala and VLPFC in conjunction with deactivation of DAN, while Network 2 engaged similar visual and limbic regions but also included distinct, adjacent DAN regions and DLPFC areas. The negative spatial component of Network 2 also comprised DMN nodes while the negative spatial component of Network 4 resembled the action mode network^41^. As such, each network is situated to process emotionally salient stimuli in conjunction with distinct prefrontally-based cognitive processes. Network 2 shows the most extensive recruitment of the occipitotemporal-VLPFC pathway, suggesting engagement of working memory processes, which might support maintenance of the task rule set, as well as dorsally mediated spatial attention which could support performance of the spatially configured matching task, particularly in conjunction with the suppression of attention to non-relevant areas of the visual field via Networks 1 and 4.

Network 6, which increased activation gradually over the course of each emotional faces block, may have a role in regulatory function. Like Network 2, Network 6 comprises the occipitotemporal-VLPFC pathway supporting object working memory as well as the occipitotemporal-amygdala pathway implicated in processing emotionally salient stimuli. It is distinguished from Network 2 by the suppression of medial visual networks, engaged during low-level processing of visual stimuli. This network may support the ongoing maintenance of rule sets and focus of visual attention as the interference load created by the emotional faces accrues across the faces block. Network 10 similarly increases in activation over the course of the faces blocks, activating most strongly withing the second half of these blocks. While still containing lateral visual areas, Network 10 primarily resembles a right-lateralized FPN component, suggesting a role for this network in top-down executive control, which become increasingly recruited over the course of the more challenging task blocks.

While the positive components of both Network 10 and Network 9 showed significant overlap with both DMN and FPN, the FPN regions of Network 9 were left-lateralized. Network 9 also dramatically differed from Network 10 in its time course, showing suppression early in faces blocks as well as early in the first shapes block, which decreased over the course of these blocks. Activation of Network 9 occurred primarily within the inter-block interval, suggesting this executive network might play a role in task switching. The time course of Network 5 closely resembled that of Network 9 despite limited spatial overlap. The suppression of Network 5, which primarily overlapped DMN, during the first shapes task block and the faces blocks likely reflects the need for increased task focus during these periods and a consequent reduction in internally-focused or self-reflective cognitive processes. The increased activation of this network during the inter-block intervals may likewise result from a temporary increase in such internal reflections.

For the Networks 3 and 7, which were rapidly activated or suppressed during the between-block transitions, activation may reflect the brief rest period, the presentation of the written instruction card, and preparation for a change in task requirements. The overlap with the language network in both cases suggests the written instructions do play a role in recruitment of these networks. The recruitment of DMN nodes and suppression of premotor and somatomotor cortex within Network 7 along with deactivation of visual networks and DAN during the intertrial interval in both networks may reflect the temporary reduction in task demands. The recruitment of the parietal memory network during these intervals in Network 3 may be due to the need to recall general task instructions. In addition to the intertrial period, these networks were also differentially engaged during the faces and shapes conditions, though the engagement of Network 3 during the first shape block resembled that observed during subsequent faces blocks and the suppression of Network 7 during shapes blocks decreased over the course of the run. These patterns converge to suggest roles for these two networks in the general recruitment and redistribution of attentional resources during the task.

Perhaps most surprisingly, despite the EFMT’s clear recruitment of multiple brain networks and regions implicated in emotion processing and internal affective state, none of the EFMT networks reflected individual differences in internalizing/negative affect or well-being/positive affect. This coincided with the absence of the subgenual cingulate, a brain region centrally linked to negative affect and depression^28,29^, as a node in any of the EFMT-recruited networks. This was particularly striking given the involvement of nearly 75% of cortex in one or more of these networks. Together with convergent findings from the systematic review of the EFMT literature^8^ and other recent studies^42,43^, our findings suggest that the EFMT may not be salient, challenging, or evocative enough to be a truly incisive tool for assessing functioning of the negative valance system.

There are some limitations to the present study. First, while the tICA identified ten task-related networks and their temporal profiles, the task performance data that are available are somewhat limited for interpreting the task components that may be engaging each network. Probing the roles of each of these networks by linking to more detailed performance measures will be an exciting new direction for research. A second limitation of our study is that the participants in the HCP study were generally healthy young adults, with no or only sub-clinical levels of anxiety, depression and drug use. This may limit our ability to link brain networks to individual variability in negative affect. However, our findings are consistent with the review by Savage and colleagues^8^, which also found no consistent relationship between brain activation in primarily emotion-processing brain areas during EFMT and mental health disorder diagnoses, including major and bipolar depression, anxiety-related disorders and obsessive compulsive disorder, among others.

In summary, we used tICA to identify and characterize spatiotemporal features of brain network processes recruited by the EFMT that have previously been missed by conventional statistical voxel-wise GLM analyses of this important task. Our findings further highlight the predominantly cognitive nature of this task and call into question the EFMT’s suitability to evoke brain activity that distinguishes individual differences in the negative valence system in health young adults.

## Methods

### Subjects

Data used in these analyses include fMRI data from the Human Connectome Project Young Adult (HCP-YA) 1200 release. A detailed description of the recruitment process for the HCP-YA is provided by others^44,45^.^44,45^. Briefly, individuals were healthy young adults and exclusion criteria included a history of major psychiatric disorder, neurological disorder, or medical disorder known to influence brain function. For the present study, we additionally excluded subjects with HCP-YA quality assessment flags, subjects with excessive head motion (i.e., 1 or more frames with framewise displacement > 2mm), and subjects without complete EFMT fMRI data. For reproducibility assessment, we randomly separated eligible subjects into two groups with non-overlapping family structure (Group 1: n=413; Group 2: n=416). Given the oversampling of siblings and twins in the HCP-YA cohort, consolidating subjects from the same family within the same group prevents family structure-related data leakage across the two groups. To maximize the generalizability of replication analyses, we did not rebalance the family-separated groups by other demographic variables (**Table 3**).

**Table 3:** Participant demographic data.

|  | **Group 1** | **Group 2** | **Test Statistic** | **p-value** |
| --- | --- | --- | --- | --- |
| **N** | 413 | 416 |  |  |
| **Gender (F/M)** | 184/229 | 262/154 | Chi-Square = 28.34 | <.001 |
| **Age (years)** | 28.21 (3.68) | 29.17 (3.75) | t(df) = 3.74 | <.001 |
| **Education (years)** | 14.93 (1.79) | 14.92 (1.78) | t(df) = 0.04 | .970 |
| mean (standard dev.) |  |  |  |  |

### fMRI Acquisition

HCP neuroimaging data were acquired with a standard 32-channel head coil on a Siemens 3T Skyra modified to achieve a maximum gradient strength of 100 mT/m^45^. T1-weighted high resolution structural images, acquired using a 3D MPRAGE sequence with 0.7mm isotropic resolution (FOV=224mm, matrix=320, 256 sagittal slices, TR=2400ms, TE=2.14ms, TI=1000ms, FA=8°), were used in the HCP minimal pre-processing pipelines to register functional MRI data to a standard brain space. Blood oxygenation level dependent (BOLD) functional magnetic resonance imaging (fMRI) data were acquired using a gradient-echo EPI sequence with the following parameters: TR = 720 ms, TE = 33.1 ms, flip angle = 52°, FOV = 280 × 180 mm, Matrix = 140 × 90, Echo spacing = 0.58 ms, BW = 2290 Hz/Px, 72 slices, 2.0 mm isotropic voxels, with a multiband acceleration factor of 8. HCP neuroimaging data were acquired in two sessions, each occurring on a different day. Two, 2-minute 16-second runs of the EFMT were acquired sequentially during session two, first with left-right (LR) phase encoding and then with right-left (RL) phase encoding.

### fMRI Preprocessing

The HCP minimal preprocessing pipeline is described in detail in Glasser et al. (2013)^46^. Briefly, minimal pre-processing of fMRI data corrects for spatial distortions, realigns volumes to compensate for subject motion, registers the fMRI data to the structural image, reduces the bias field, normalizes the 4D image to a global mean, and masks the data with a final FreeSurfer-generated brain mask^46^. We additionally applied 4mm spatial smoothing, 150 sec temporal smoothing, removal of non-steady-state frames (first 15 frames), and ICA-FIX denoising;^47^ which denoises residual subject motion, physiological motions, and artifacts, to the HCP minimally preprocessed EFMT fMRI data.

### Emotional Face Matching Task (EFMT)

The EFMT, as implemented for HCP-YA, was adapted from Hariri and colleagues^2^. Subjects were presented with alternating blocks of trials that asked them to decide either 1) which of two faces presented on the bottom of a screen matched a face at the top of the screen, or 2) which of two shapes presented at the bottom of a screen matched the shape at the top of the screen (shapes were ovals, participants matched orientations). The faces all have either angry or fearful expressions. Each of the two EFMT runs included 3 face blocks and 3 shape blocks. Each block included 6 trials of the same stimulus type (i.e., faces or shapes), wherein each trial consisted of a 2 second stimulus presentation followed by a 1 second intertrial interval (ITI). Each block was also preceded by a 3 second task cue (“face” or “shape”), making each block 21 seconds in total.

Given a strong ceiling effect in matching accuracy on this task, we prioritized reaction times in assessing task performance. More specifically, to gauge individual differences in emotion interference (i.e., the extent to which processing emotional stimuli slows subjects’ ability to complete the task matching goal), we targeted the difference in reaction time between face-matching trials and shape-matching trials as our primary task performance metric.

### fMRI Analysis

We applied tICA to four separate runs of FIX-denoised EFMT fMRI data: Group 1 LR, Group 1 RL, Group 2 LR, and Group 2 RL. This identified task-relevant brain networks, including their voxel-wise spatial maps and their frame-wise time courses, and the individual subject loadings on each network, for each run in both groups. For Group 1 LR, we ran tICA at varying model orders between d=5 and d=25 to inspect the output for the optimal model order for analysis, which we identified to be d=20 in line with previous reports examining the correspondence between task activation networks and canonical resting state brain networks^48,49^. The time courses, power spectra and spatial maps of these 20 components were then examined to identify those most likely to reflect task-related brain activity (e.g., components with spatial patterns resembling large-scale brain networks who had temporal courses reflecting different components of task performance). The identified network components were assessed for both spatial and temporal reproducibility across the LR and RL acquisitions and across Group 1 and Group 2, with Group 1 LR as the reference data. We then applied hierarchical clustering to the time course correlation matrix of identified EFMT-network components from Group 1 LR to distinguish groups of networks with overlapping and distinct time courses.

To examine correspondence between network time courses identified by tICA and the task stimulation time course (illustrated in **Fig. S4**), the ten network time courses were fit *post hoc* with a standard block design GLM with four regressors: face blocks and shapes blocks -each convolved with a gamma hemodynamic response function - and the temporal derivatives of each. A Bonferroni-corrected threshold of p < .0005 was applied to account for ten networks and ten contrasts of interest.

To further characterize the spatial maps of the ten task networks, we used the network correspondence toolbox (NCT)^31^ to compute overlap (Dice coefficients) between the positive and negative components of each task network and each of 17 canonical large-scale networks defined in the Gordon et al. parcellation^30^. In order to assess statistical significance, a spin test implemented in Python was performed with 1000 permutations for each reference atlas network^31^. Significant overlap was thresholded at p<0.05.

## Notes

### Competing Interest Statement

The authors have declared no competing interest.

