## Supplemental Material for "Concurrent large-scale brain dynamics during the emotional face matching task and their relation to behavior and mental health"

Supplementary Materials

**Figure S1.** Hierarchical clustering of EFMT-network timeseries.

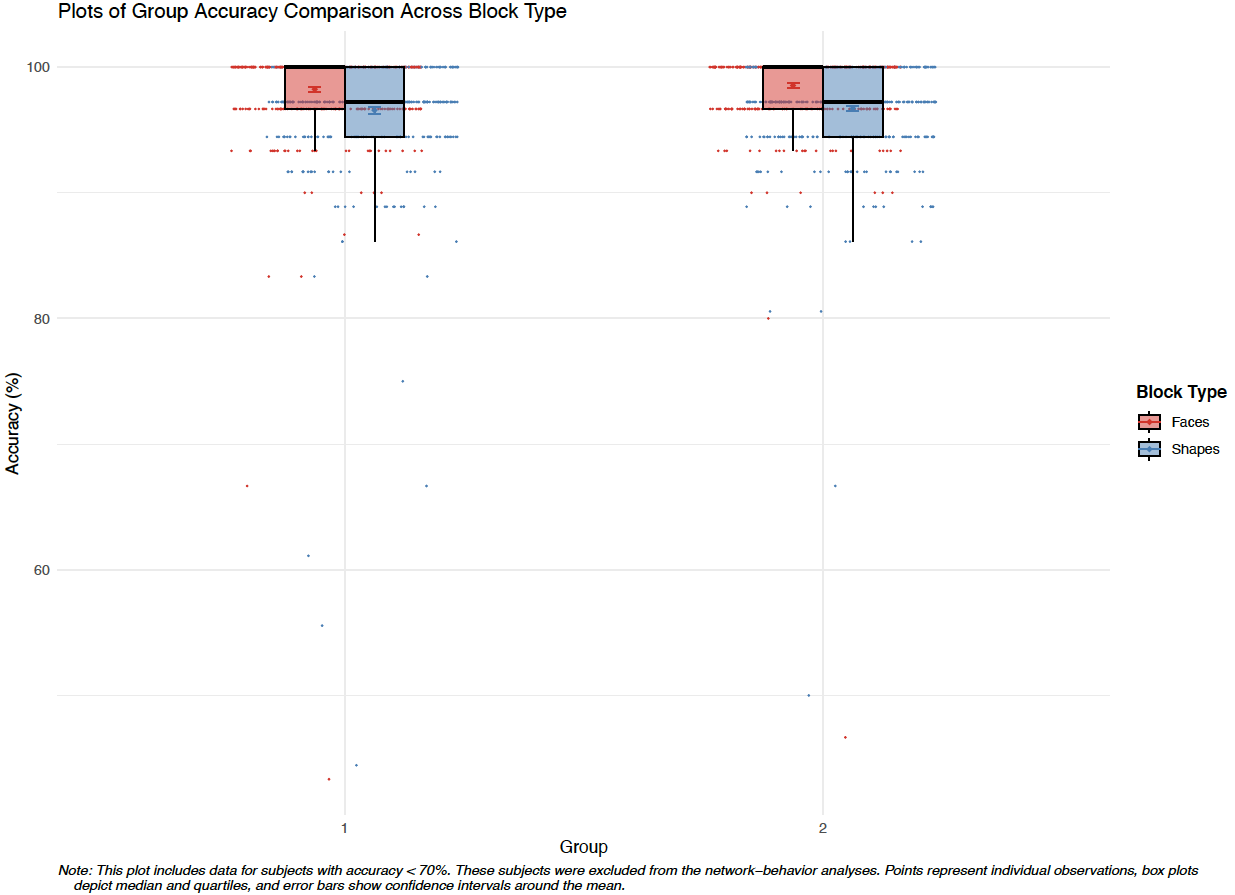

**Figure S2.** Group average accuracy on faces blocks and shapes blocks for Groups 1 and 2. Points represent individual observations.

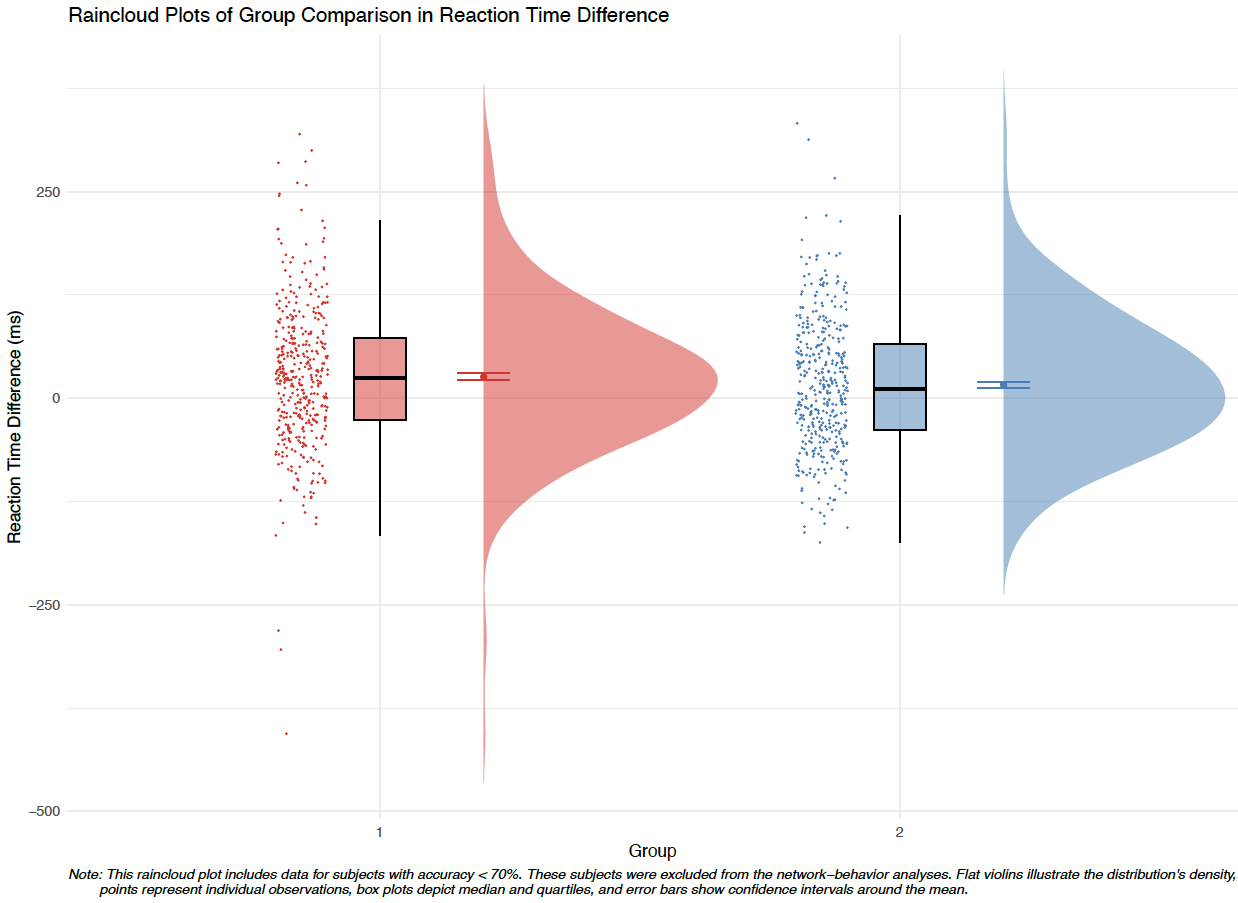

**Figure S3.** Group average “emotion interference” for Groups 1 and 2.

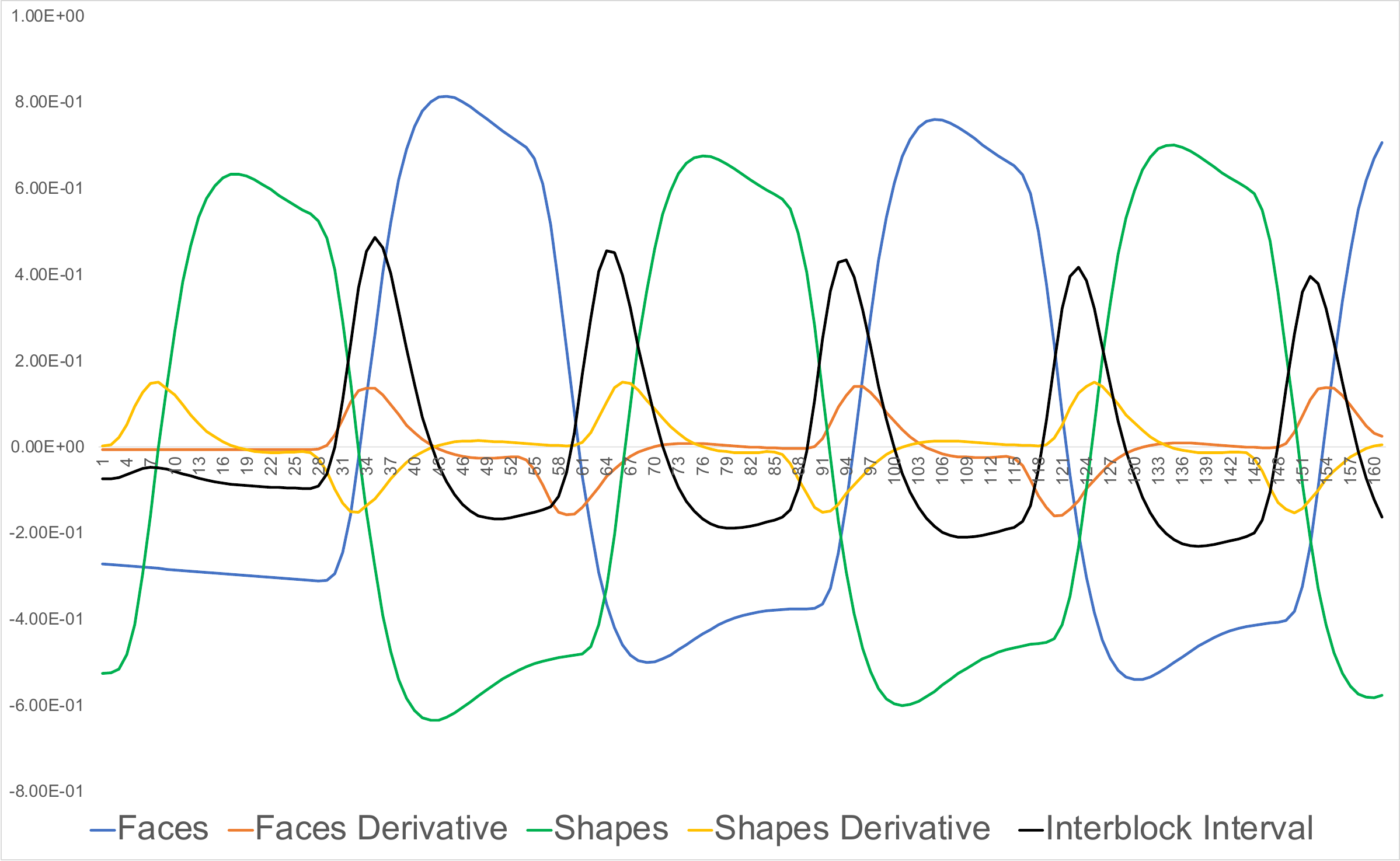

**Figure S4.** EFMT contrast time courses.

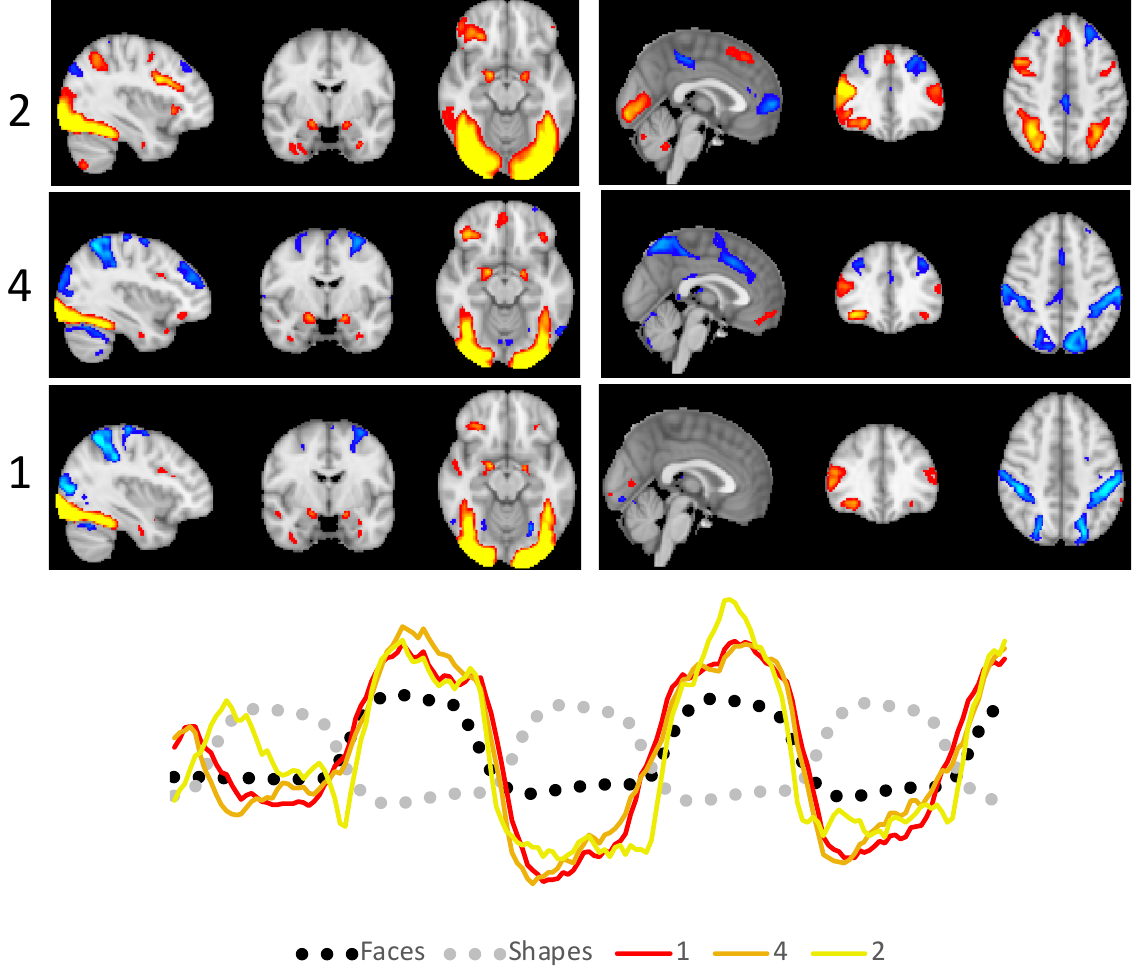

**Figure S5.** Spatial and temporal comparison of Networks 1, 2 and 4.

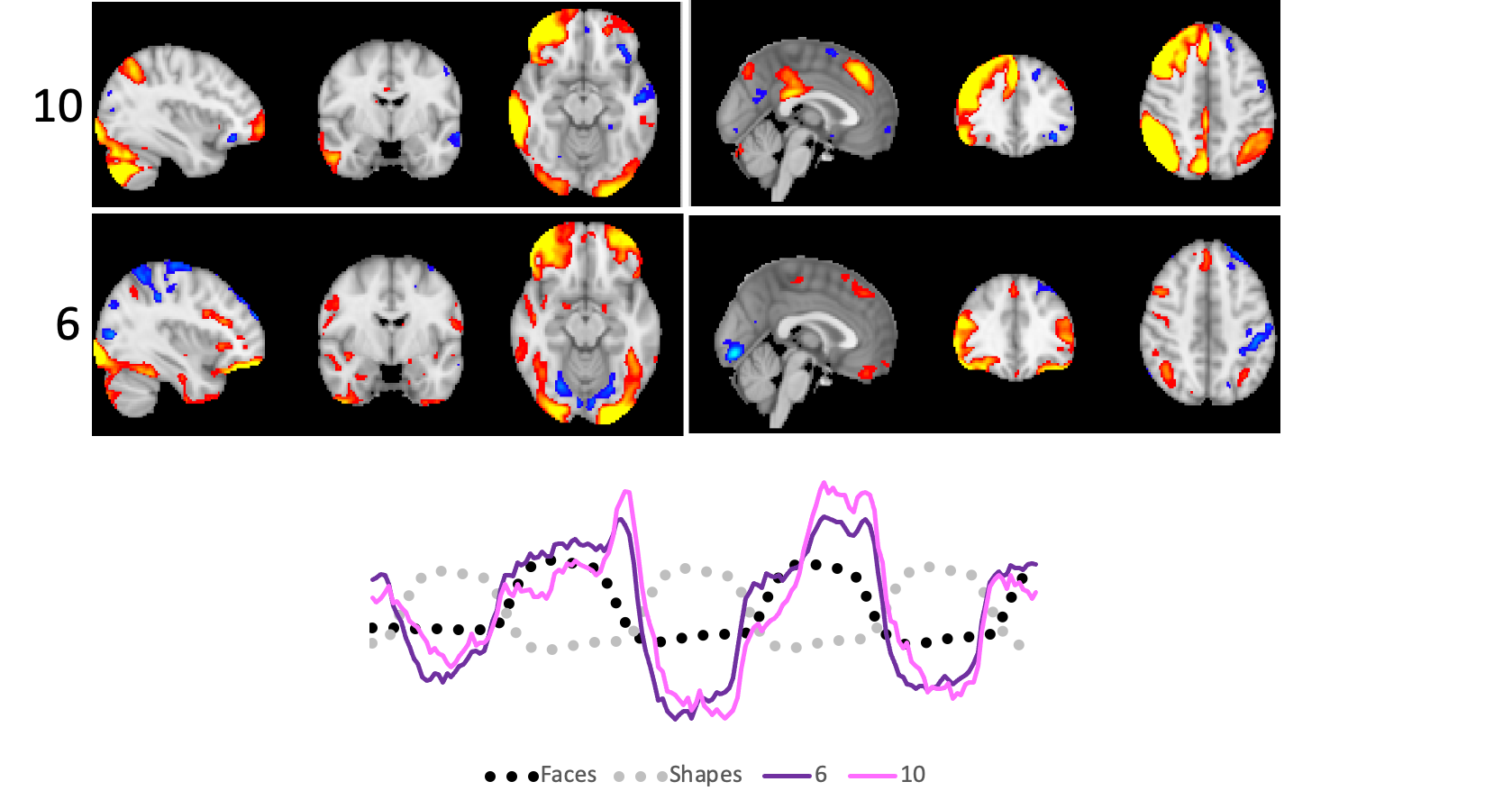

**Figure S6.** Spatial and temporal comparison of Networks 6 and 10.

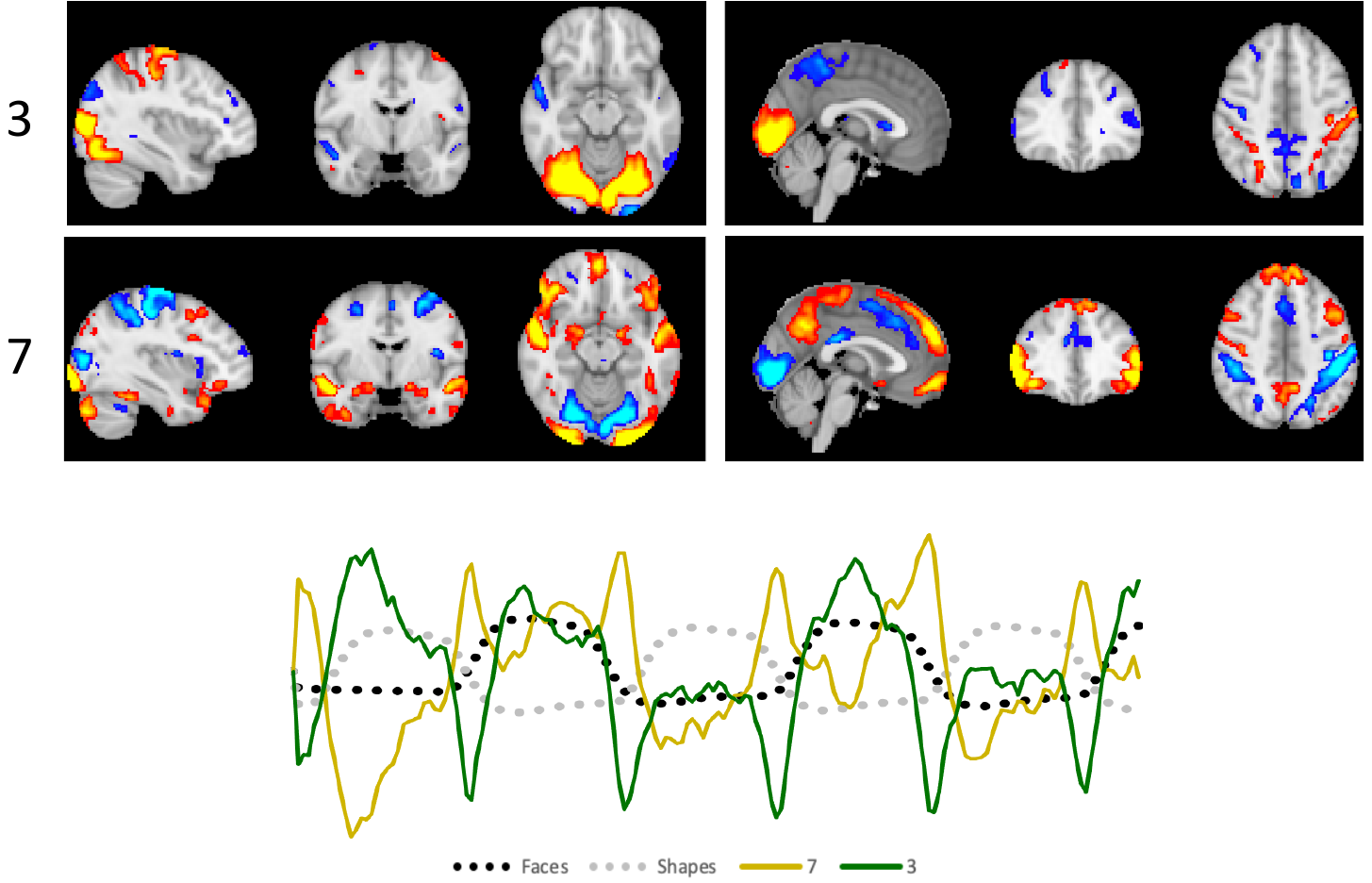

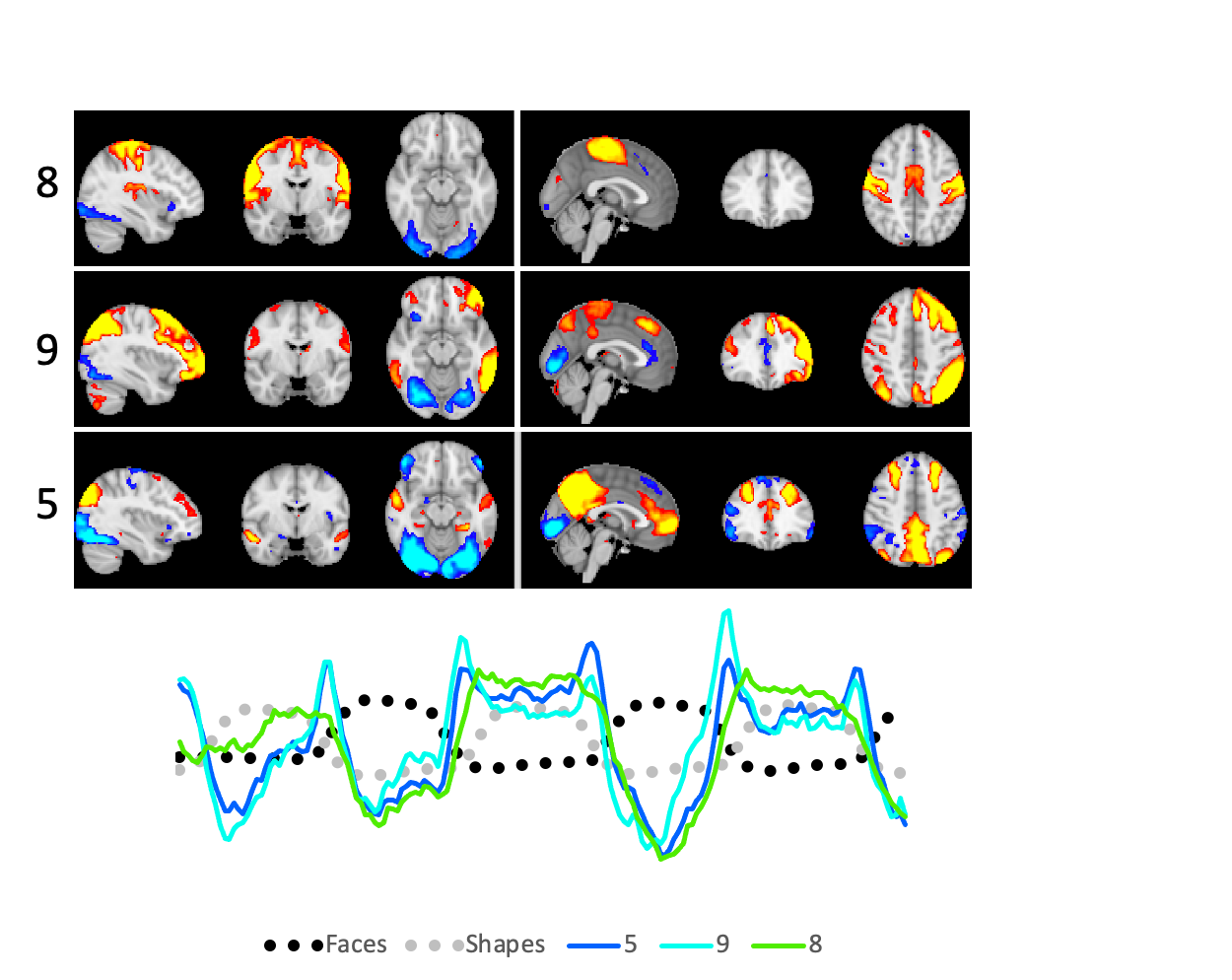

**Figure S7.** Spatial and temporal comparison of Networks 3 and 7.

**Figure S8.** Spatial and temporal comparison of Networks 5, 8 and 9.
